# Application of the radial distribution function for quantitative analysis of neuropil microstructure in stratum radiatum of CA1 region in hippocampus

**DOI:** 10.1101/003863

**Authors:** Yuriy Mishchenko

## Abstract

Various structures in the brain contain many important clues to the brain?s development and function. Among these, the microscopic organization of neural tissue is of particular interest since such organization has direct potential to affect the formation of local synaptic connectivity between axons and dendrites, serving as an important factor affecting the brain?s development. While the organization of the brain at large and intermediate scales had been well studied, the organization of neural tissue at micrometer scales remains largely unknown. In particular, at present it is not known what specific structures exist in neuropil at micrometer scales, what effect such structures have on formation of synaptic connectivity, and what processes shape the micrometer-scale organization of neuropil. In this work, we present an analysis of recent electron microscopy reconstructions of blocks of neuropil tissue from rat s. radiatum of hippocampal CA1 to provide insights into these questions. We propose a new statistical method for systematically analyzing the small-scale organization of neuropil based on an adaptation of the approach of radial distribution functions from statistical physics. Our results show that the micrometer-scale organization of hippocampal CA1 neuropil can be viewed as a disordered arrangement of axonal and dendritic processes without significant small-scale positional coordinations. We observe several deviations from this picture in the distributions of glia and dendritic spines. Finally, we study the relationship between local synaptic connectivity and the small-scale organization of neuropil.

## Introduction

Understanding the brain’s anatomical structure is an important goal of neuroscience both from the point of view of producing new insights into the principles of brain’s organization (Arbib et al., 1997; Bono and Villu Maricq, 2005; Hatton, 1990; Sporns et al., 2004, 2005) and better understanding of the brain’s development and disorders (Brambilla et al., 2003; Castellanos et al., 2002; Garrard et al., 1998; Geschwind, 1975; Good et al., 2002; Uhlhaas and Singer, 2006). Diverse anatomical structures are readily observable in the brain at a variety of scales. At the largest scale, the organization of the brain into specific regions and functional areas has been well established (Brodmann and Garey, 2005; Damasio, 2005; Kandel et al., 2000). At intermediate scales, structures such as cortical layers (Kandel et al., 2000; Nolte, 2002), cortical columns (Freeman, 2003), topographic mappings (Adams and Horton, 2003; Montero et al., 1977), ocular dominance patterns (Erwin et al., 1995; Miller et al., 1989) and orientation selectivity patterns in visual cortex (Bosking et al., 1997; Ohki et al., 2005), as well as topographic mappings in auditory (Morosan et al., 2001; Pickles, 2012) and somatosensory cortex (Nelson, 2001) are also known. A range of genetic, biomolecular, and neural activity mechanisms are known to be associated with the formation and development of these structures, providing important clues about the brain’s development and dysfunctions (Dickson, 2002; Ferster and Miller, 2000; Keil et al., 2010; McLaughlin and O’Leary, 2005; Parrish et al., 2007; Tear, 1999; Wolf et al., 2011; Wong, 1999).

Despite this extensive body of knowledge, the anatomical organization of the brain at micrometer scales - at the level of neural tissue - remains largely unknown. Such organization is of general interest in neuroscience. It is also of substantial practical significance. It had been hypothesized that neuropil’s small-scale organization can immediately affect the formation of synaptic connectivity between axonal and dendritic processes in neuropil, via availability of local axonal and dendritic partners for synaptic connections (Braitenberg and Schuz, 1998; Peters, 1979; Stepanyants and Chklovskii, 2005; Stepanyants et al., 2008). Abnormal changes in such small-scale neural tissue organization, therefore, can be an important factor contributing to the development of neurodegenerative disorders, frequently characterized by degeneration of synaptic connectivity in neural tissues (Bonda et al., 2010; Hamos et al., 1989; Raff et al., 2002; Scheff et al., 2006; Terry, 2000).

In the present study, we attempt to new yield insights into micrometer-scale organization of neural tissue using recent anatomical serial section Transmission Electron Microscopy (ssTEM) reconstructions of several blocks of neuropil from stratum radiatum of hippocampal CA1 tissue in rat (Mishchenko, 2009; Mishchenko et al., 2010). To address this question, we propose a new quantitative approach based on an adaptation of the statistical tool of radial distribution function from material sciences (Chandler, 1987; McQuarrie, 2000; Sandler, 2010). Radial distribution function is a statistical tool that can be used to characterize spatial correlations existing between different structural elements of a material. Radial distribution functions have been used commonly in statistical physics to analyze the organization of various materials including complex alloys, solutions, and colloids.

We perform a direct measurements of radial distribution functions for hippocampal CA1 neuropil using the neuropil tissue reconstructions from (Mishchenko et al., 2010). We analyze obtained measurements using statistical and modeling approaches. The results of our analysis indicate that the organization of neuropil at micrometer scales most closely resembles a disordered arrangement of neural processes, spines, and glia fragments. We also discuss the deviations from this picture observed in the distributions of dendritic spines and glia in neuropil. Finally, we analyze the variations in the spine density of different dendritic segments in relation to the structure of surrounding neuropil, as measured by the radial distribution function, and find that no significant correlation exists between these two variables.

To summarize, in this work we produce new results about the organization of neural tissue in the brain at the scales of 1-10 μm. We analyze several blocks of rat hippocampal CA1 tissue, reconstructed using ssTEM. We use a new quantitative approach for the analysis of microscopic neuropil organization based on an adaptation of radial distribution functions from material science, in order to determine presence of any “nonrandom” structures in neuropil’s organization at micrometer scales. Our results show that the organization of CA1 neuropil at micrometer scales is essentially disordered or stochastic, at least to the extent characterized by the radial distribution functions we measure.

## Materials and Methods

### Neuropil data

Three approximately cubic blocks of hippocampal neuropil tissue reconstructed using ssTEM in (Mishchenko et al., 2010) were used to perform the analysis in this work. The datasets 1, 3, and 4 from in (Mishchenko et al., 2010), specifically, were used; the dataset 2 was not used because of its small size - a 30 μm^3^ neuropil block surrounding a single dendritic spine. All volumes were from the middle of s. radiatum about 150 to 200 micrometers from the hippocampal CA1 pyramidal cell layer. Volumes 1 and 3 were from a perfusion-fixed male rat of the Long-Evans strain weighing 310 gm (postnatal day 77) as described in (Harris and Stevens, 1989)), and volume 4 was from a hippocampal slice from a postnatal day 21 male rat of the Long-Evans strain fixed as described in (Fiala et al., 2003). All animal procedures were performed in accord with relevant regulatory standards, as detailed in (Fiala et al., 2003; Harris and Stevens, 1989). To produce the said datasets, briefly, the series of ultrathin sections were cut at ~45-50 nm using an ultramicrotome, mounted and counter stained with saturated ethanolic uranyl acetate followed by Reynolds lead citrate. Sections were then photographed on JEOL 1200EX and 1230 electron microscope (JEOL, Peabody, MA) at magnification of 10,000X or 5,000X, and stored as series of digital images at 4.4 nm/pixel. The neuropil contents of these images were reconstructed using the approach of (Mishchenko, 2009) in Matlab and stored as Matlab data ready for further analysis. The reconstructions totaled 670 μm^3^ and contained 1900 fragments of different neural processes (Figure 1).

**Figure 1:**
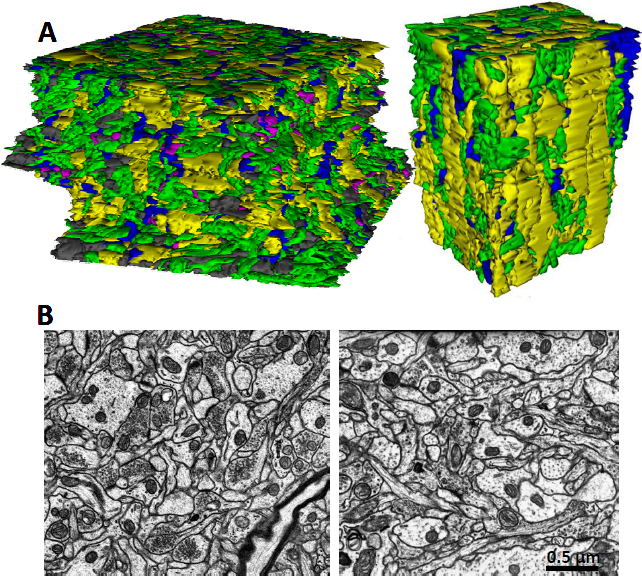
Dense ssTEM reconstructions of hippocampal CA1 neural tissue reveal complex organization of neuropil at micrometer scales. (A) An example of dense 3D ssTEM reconstruction “Volume 1” from (Mishchenko et al., 2010) (left) and “Volume 4” from (Mishchenko, 2009) (right). The respective neuropil blocks measure 9.1 × 9.0 × 4.1 μm^3^ and 6.0 × 4.3 × 5.1 μm^3^. Different colors represent the neural processes of different types: green for axons, yellow for dendrites, magenta for spines, and blue for glia. Gray represents the objects that could not be classified based on ssTEM images alone. (B) An example of ssTEM micrograph images from “Volume 1” (left) and “Volume 4” (right).

### Definition of radial distribution functions

Radial distribution functions are defined conventionally in statistical physics as the change in the density of the particles of a certain sort in concentric radial shells surrounding a typical “reference” particle in a material (Figure 2A). Radial distribution functions characterize spatial correlations present in a material and, in that capacity, provide a powerful tool for systematic analysis of the small-scale organization of complex materials and systems. In material sciences, radial distribution functions have been used commonly to analyze the organization of different complex materials such as alloys, solutions, colloids, etc.

**Figure 2:**
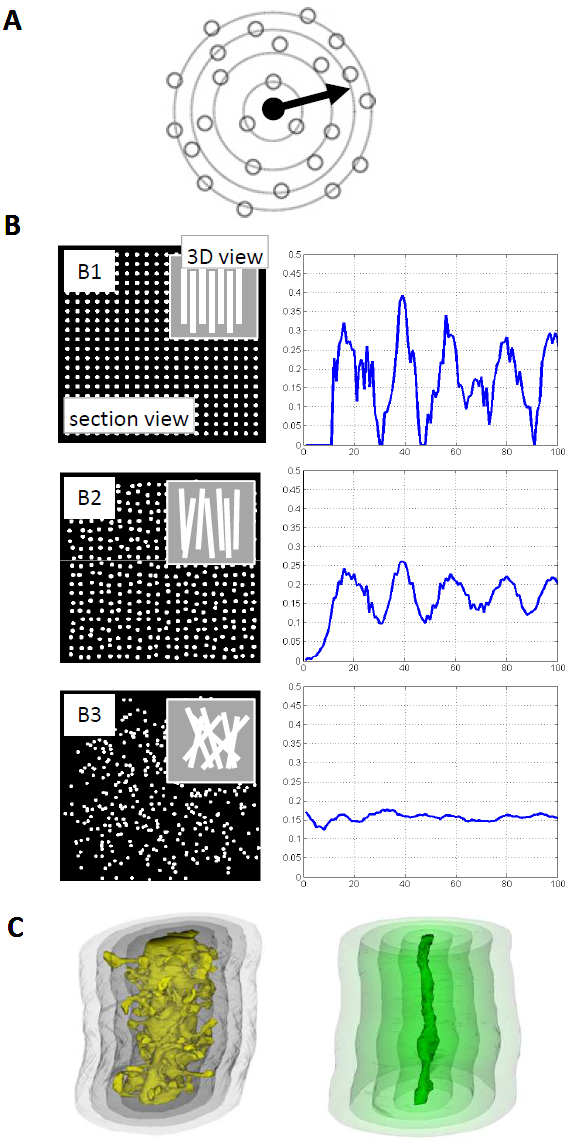
Radial distribution functions as the tools for characterizing small-scale structure of neuropil. (A) Radial distribution functions in physical material science offer a powerful tool for characterization of the small-scale structure of complex materials and systems. Radial distribution functions are defined as the change in the density of the particles of a certain sort in concentric shells surrounding a “typical” reference particle in material. Radial distribution functions can be visualized as counting the number of the particles in each concentric shell around a fixed reference particle, divided by the volume of that shell. (B) An example of radial distribution functions calculated for different idealized neural processes’ arrangements in neuropil. Panel B1 shows an arrangement that is perfectly ordered, that is, such where all neural processes are oriented in the same direction and are also spatially aligned (left). The radial distribution function in this case shows strong peaks representative of the spatial correlations present in this arrangement (right). Panel B2 shows an arrangement where neural processes are partially disordered (left). In this arrangement, the radial distribution function still possesses peak-like features, but these become less pronounced due to the destruction of correlations (right). Panel B3 shows an arrangement that is completely disordered (left). The radial distribution function in this case is flat and lacks any significant features (right). (C) In the case of neuropil, due to the irregular one-dimensional nature of neural processes, we modify the conventional definition of radial distribution function to use cylindrical concentric shells constructed relative to the surface of axonal or dendritic processes. The calculation of such radial distribution function is exemplified here using a series of cylindrical concentric shells constructed at different distances away from the surface of one reference dendrite (left) or one reference axon (right). Radial distribution functions for neuropil are defined as the average density of neural processes of different kind in such shells at different distances away from “typical” reference axons or dendrites.

Radial distribution functions can be visualized easily by counting the number of particles present in each concentric spherical shell surrounding a fixed reference particle. If particles in a system are distributed completely randomly and uniformly, then such counts would grow in direct proportionality to the volume of the shells, whereas their density in each shell would be a constant. In other words, the radial distribution function in such a situation is simply a constant. If a system possessed strong interactions between particles leading to them having different likelihoods of appearing at different relative distances, then such density would vary from shell to shell, resulting in a radial distribution function with peaks and other distinctive features. In general, we can say that in disordered materials the radial distribution functions are featureless and flat, while in ordered systems the radial distribution functions possess peaks and other distinctive features, representative of the spatial correlations present in such systems.

Figure 2B shows examples of radial distribution functions calculated for different simulated idealized arrangements of neural processes in a neuropil. Panel B1 shows an example of such an arrangement that is perfectly ordered, that is, such where all neural processes are oriented in the same direction and are also spatially aligned. The radial distribution function for this case is shown in the right subpanel: the respective radial distribution function shows strong peaks representative of the spatial correlations present in this arrangement. Panel B2 shows a model arrangement where neural processes are partially disordered, that is, such where their positions are less well coordinated and also their headings may vary slightly. In this arrangement, the radial distribution function still possesses peak-like features, but such peaks become smoother and less pronounced due to the destruction of the correlations. Finally, panel B3 shows an example of such an arrangement that is completely disordered, that is, where the headings of neural processes are completely random as are their positions. The radial distribution function in this case is flat and shows no significant features.

As had been discussed above, radial distribution functions are defined conventionally using concentric spherical shells, in order to quantify the changes in system components’ densities with the distance from a point-like reference particle. In the case of neuropil, however, we adopt a slightly different definition because of the irregular cross-section shapes of axonal and dendritic processes and also their elongated fiber-like nature. Specifically, we define the radial distribution functions in neuropil as the average density of different neural processes, namely axons, dendrites, spines, and glia, in *cylindrical* concentric shells constructed relative to the surface of the respective reference neural process, Figure 2C.

It shall be noted that radial distribution functions, as a signature of organization of a material or a system, are sensitive to a particular type of structures such as positional coordination, that is, the situations where the presence of one neural process influences the likelihood of encountering neural processes of other types in its vicinity. Thus, while radial distribution function is highly informative about spatial organization of a material, it is not be sensitive to all possible structures, especially such that do not manifest immediately in spatial position or that depend on non-spatial properties such as excitatory/inhibitory nature of neural process, etc.

### Measurement of radial distribution functions in neuropil

In order to calculate the radial distribution functions in hippocampal neuropil, first we constructed a series of concentric equidistant cylindrical shells from the surface of each axon and dendrite in our dataset (approximately 1500 neural processes in total). This was done by calculating Euclidian distance transform (EDT) for neural process. Distance transform is an operation on the dataset that assigns to every pixel in the volume the shortest distance from that pixel to the surface of a respective object, here, a neuronal process. This calculation was performed using a custom implementation of EDT algorithm for very-large-size datasets in Matlab described in (Mishchenko, 2013). The set of equidistant surfaces of such distance transforms constituted a set of concentric cylindrical shells that could be used for calculating the radial distribution functions of interest (see Figure 2C).

A set of radial distribution functions for each neural process was calculated by computing the density of different neuropil components in the above concentric shells around each neural process at the step of 13.2 nm in volumes 1 and 3 and 8 nm in volume 4. The latter difference was due to the different pixel size in volume 4 (8 nm/pixel vs. 4.4nm/pixel).

Thus produced individual radial distribution functions were averaged over all reference axons and dendrites to produce the respective neuropil’s average axon- and dendrite-specific radial distribution functions.

In the case of dendrites, the above procedure was carried out relative to dendritic shafts of the dendrites by first applying an algorithm that truncated dendritic spines at the bases of their necks. All dendritic shafts were subsequently manually examined to insure that no attached spines remained, which would have the result of damaging the result of measuring the respective radial distribution functions.

### Measurement of the size distribution of axonal and dendritic cross-sections in neuropil

In order to measure the distribution of the local sizes of axonal and dendritic cross-sections in hippocampal neuropil, we first evaluated 3D distance transform inwards of each neural process in the sample. This had the effect of assigning to each pixel interior to a neural process the shortest distance from that pixel to the containing process’s surface. The centerline of each neural process was calculated as the set of points where such distance was a local maximum in at least two out of three orthogonal directions. The values of the distances were then collected along such centerlines for each neural process, characterizing the size distribution of the diameters of their cross-sections at a step of approximately 50 nm. The class-specific distributions were calculated by averaging such distributions over all axonal and dendritic processes, respectively.

### The model of random mixing of local neuropil

To quantitatively describe the salient features of measured neuropil radial distribution functions, we propose a local random mixing model for neuropil. The aim of this model is to describe how the density of axonal and dendritic processes changes in concentric shells because of termination and re-appearance of axonal and dendritic cross-section profiles in the shells as such shells sweep through neuropil volume.

If we imagine ourselves moving through the concentric radial shells away from the surface of one reference axon or dendrite, it is easy to imagine that the cross-sections of other axons and dendrites would emerge, grow, contract, and disappear as such shells pass through different axonal and dendritic processes in neuropil. Consequently, the volume occupied by axonal and dendritic cross-sections in each shell would change as the result of these processes of emergence and disappearance of axonal and dendritic profiles. In random mixing model, we model these processes using a constant rate that corresponds to the rate with which such processes occur on average in the sample’s volume. For example, if we imagine a large surface moving through a volume with uniformly mixed axonal and dendritic processes, axonal and dendritic cross-sections would appear and disappear with a frequency close to the inverse of the respective processes’ diameters. In other words, as cylindrical radial shells sweep through neuropil volume, axons appear and disappear in the shells frequently due to their small crosssection size, while dendritic cross-sections appear and remain in the shells for much longer periods of time. If such cross-sections were to appear and disappear in the radial shells completely randomly, with defined rates, then the radial distribution function would be imparted a particular shape, which we calculate below.

To formalize the above intuition mathematically, we consider the process of moving away from the surface of a reference axon or dendrite over a set of concentric radial shells. If the average diameters of axonal and dendritic processes in the sample volume are given by *d_axn_* and *d_dnd_*, the probability of an axon or a dendrite cross-section terminating in the shell as one moves over a distance Δ*x* can be approximately described as Δ*x/d_axn_* and Δ*x/d_dnd_*. The space vacated in the shell due to such terminations would be subsequently re-partitioned among newly emerged axons and dendrites, which occurs in the proportion of the total surface areas of axonal and dendritic processes in the volume, *n_axn_*π*d_axn_L* : *n_dnd_*π*d_dnd_L*. Here, *L* is a constant defining the linear size of the sample volume and *n_axn_*, *n_dnd_* are the number densities of axons and dendrites in the volume, respectively, whereas *πdL* is the surface area of one neural process viewed as a straight cylinder of diameter *d* and length *L*.

We denote the relative densities occupied by axonal and dendritic processes in cylindrical radial shell at distance *x* away from the reference axon or dendrite as *N_axn_*(*x*) and *N_dnd_*(*x*), respectively. Then, the above processes can be mathematically represented by two equations for the evolution of *N_axn_*(*x*) and *N_dnd_*(*x*) with *x*,

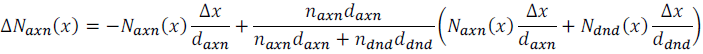

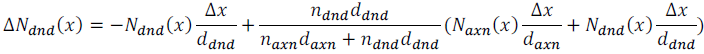

These equations describe the change in the volume occupied in the shell by different axonal and dendritic cross-sections over a step of size Δ*x*. The first term represents the termination of axons and dendrites with frequency Δ*x/d_axn_* and Δ*x/d_dnd_*, and the second term represents the re-partitioning of thus vacated space among newly emerged axons and dendrites in the ratio of axonal and dendritic surface areas, *n_axn_d_axn_* : *n_dnd_d_dnd_*. Note also that *N*(*x*) are relative densities, that is, *N_axn_*(*x*) + *N_dnd_*(*x*) = 1 at all times.

The above equations are a system of simple finite-difference equations that can be solved using standard methods. The solutions yield several immediate predictions for the respective radial distribution functions. Specifically, at small distances, *x*≈0, we find that the radial distribution functions are in the ratio equal to that of the total surface areas of dendritic and axonal processes in the sample, *N_axn_*(*x* ≈ 0):*N_dnd_*(*x* ≈ 0)≈ *n_axn_d_axn_*: *n_dnd_d_dnd_* (this can be seen, for example, by taking *N_axn_*(0) = 1 or *N_dnd_*(*x*) = 1 and performing a single step Δ*x*). At large distances, *x* ⟶ ∞, the values of the radial distribution functions are asymptotically defined by the ratio of the average volume densities of axonal and dendritic processes in the sample, *N_axn_*(∞):*N_dnd_*(∞) ≈ *n_axn_d_axn_*^2^: *n_dnd_d_dnd_*^2^ (this can be seen by enforcing in the above equations the condition Δ*N_axn_*(∞) = Δ*N_dnd_*(∞) = 0). Given *d_dnd_* > *d_axn_*, it follows that these ratios always differ by the factor of *d_dnd_/d_axn_.* Referring to Table 3 in (Mishchenko et al., 2010) we obtain *d_dnd_/d_axn_*≈3. Finally, the transition between the small distance and the large distance regimes occurs exponentially on the scale 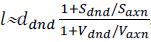, where *S_dnd_*/*S_axn_* and *V_dnd_*/*V_axn_* are the ratios of dendritic and axonal surface and volume fractions in the samples, respectively. Referring to Figure 2 in (Mishchenko et al., 2010), we obtain 1≈500 nm. Moreover, the radial distribution functions are the same whether the reference object is an axon or a dendrite, that is, the dendrite- and axon-specific radial distribution functions cannot differ. Inspection of Figure 3 in Results shows that these predictions are in excellent agreement with the radial distribution functions observed in this work.

**Figure 3:**
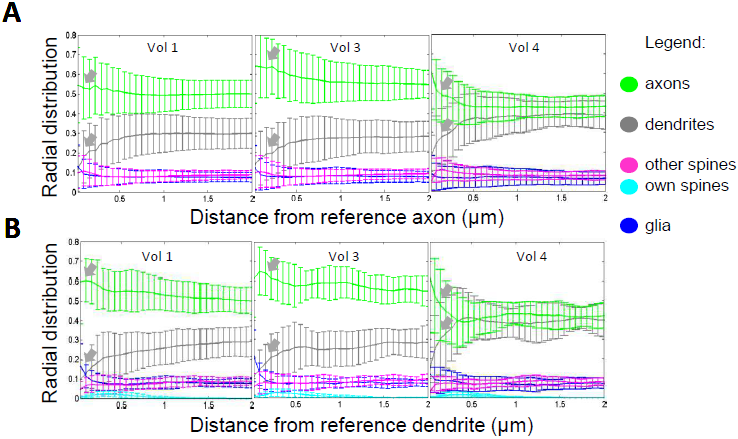
The radial distribution functions of hippocampal CA1 neuropil. (A) Radial distribution functions measured for different neuropil components relative to axonal reference processes in sample volumes 1, 3 and 4. (B) Radial distribution functions measured for different neuropil components relative to dendritic reference processes in sample volumes 1, 3 and 4. The measured radial distribution functions show predominantly flat behavior, indicating lack of significant spatial correlations among neuropil components at the scales of one to several micrometers. The error bars represent the variation in the radial distribution functions measured around different axonal and dendritic references in the samples.

## Results

### 1. Organization of hippocampal CA1 neuropil at micrometer scales

Reconstruction of complete volumes of neuropil tissue had been recently produced using electron microscopy. A quick inspection of these reconstructions reveals complex small-scale organization of hippocampal neuropil without apparent patterns. However, one cannot immediately conclude whether this lack of features is due to the absence of any structures in neuropil at micrometer scales or because such structures are complex and simply are visually obscured.

In order to provide an answer to this question, we perform a measurements of radial distribution functions for hippocampal neuropil using the reconstruction of blocks of rat s. radiatum hippocampal CA1 tissue from (Mishchenko et al., 2010), Figure 3. Please refer to Materials and Methods for the discussion of the definition and the properties of such radial distribution functions. Here and below, we will also refer to such radial distribution functions specifically in neuropil as the neuropil’s *structure functions*.

The main feature of the measured neuropil structure functions observed in Figure 3 is their flat behavior at the distances above of approximately 0.5 μm. A linear regression to structure functions at the range of 0.5-2 μm gives the coefficient of regression R=0.019±0.030 μm^−1^ with residual mean-square-error (MSE) MSE=0.0043±0.0022 μm^−1^ for dendritic structure functions, and R=−0.014±0.032 μm^−1^ and MSE=0.0035±0.0015 μm^−1^ for axonal structure functions. Flat behavior of the structure functions in this region is representative of the absence of correlations in the relative position of different neural processes in the neuropil samples at distances above 0.5 μm.

The measured structure functions in Figure 3 show peak/drop-like features at the distances of 0-0.5 μm, indicating presence of correlations between neural processes at these small distances (gray arrows in Figure 3). These features can be understood as the effect of physical contact interactions between adjacent neural processes in densely packed neuropil. That is, if at large distances away from a reference process the neural processes can mix uniformly, near the surface of that process the presence of the process’s boundary causes adjacent neural processes to re-align along it and induces correlations in the positions of such processes, reflected by the above mentioned peak/drop-like features.

This effect can be described more quantitatively using a model of random mixing of neuropil (Materials and Methods), which can reproduce well the main properties of the observed structure functions. In particular, this model predicts that near the surface of the reference object the ratio of the axonal and dendritic structure functions should equal the ratio of axonal and dendritic surface areas in neuropil. This ratio can be calculated using the data in (Mishchenko et al., 2010) as 6:1 for volumes 1 and 3 and 3:1 for volume 4, in good agreement with the structure functions in Figure 3. Furthermore, the random mixing model prescribes that at large distances the ratio of axonal and dendritic structure functions converges to the ratio of the axonal and dendritic volume densities, which can be calculated from (Mishchenko et al., 2010) as 2:1 for volumes 1 and 3 and 1:1 for volume 4, again in good agreement with Figure 3. The transition between the two regimes occurs exponentially on the length scale of *l*≈500 nm, as defined primarily by the average dendritic and axonal diameters, also in agreement with Figure 3. Finally, such the structure functions are predicted to be the same for axonal and dendritic references, again in good agreement with the measurements.

Certain differences can be noted between the structure functions measured in volumes 1 and 3 and that in volume 4. Such differences can be attributed to volume 4 being slice preparation and volumes 1 and 3 being perfusion fixed, or to that volume 4 is from a young (P21) and volume 1 and 3 are from a mature (P77) rat. Unfortunately, the small size of available data prevents us from gaining a better understanding of these differences.

Thus, we arrive at conclusion that the organization of hippocampal CA1 neuropil at the scales of one to several micrometers, as quantified by the neuropil structure functions, resembles a space-filling arrangement of axonal and dendritic processes without local order or spatial structures.

### 2. Orientational organization of hippocampal CA1 neuropil at micrometer scales

Even though the micrometer-scale organization of neural processes in neuropil may not exhibit spatial local ordering, it can still exhibit a global order such as directional alignment. In fact, it is well known from past anatomical studies that axonal and dendritic processes in hippocampus are running in approximately perpendicular directions (Amaral and Witter, 1989; Cajal, 1909; Schaeffer, 1892; Westrum and Blackstad, 1962). Here, we measure the distribution of the headings of dendritic and axonal processes in our samples. Indeed, we find that the approximately perpendicular organization of dendrites and axons is observed at these very small scales, Figure 4. We observe that dendrites tend to run collinearly approximately in the direction orthogonal to the CA1 cell-body layer and axons tend to pass mostly within the plane of cell-body layer. The axonal component is also observed to possess an interesting two-component structure, with approximately half of all axons traveling in a singly collinear bundle with common direction and the rest appearing to traverse the sample diffusely in all directions within the plane parallel to the cell-body layer. This division of axons into 50% “perpendicular” and 50% “random” components is interesting. However, we were unable to identify any significant correlates between the axonal properties (for example, axonal diameter) and this division. A more detailed study in the future may help shed new light onto this peculiar feature of microscopic axonal organization in CA1 neuropil.

**Figure 4:**
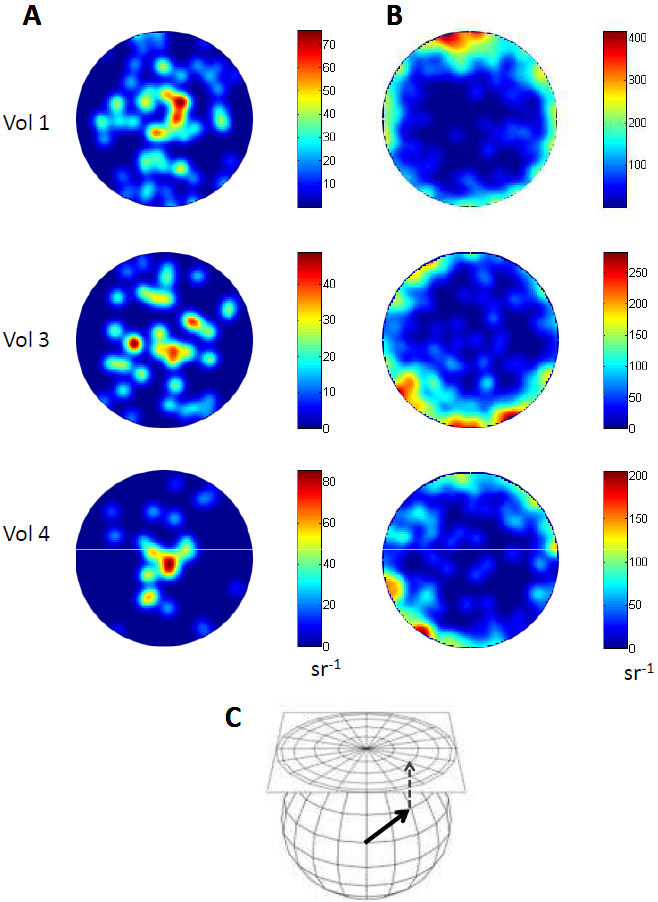
The directional organization of hippocampal CA1 neuropil. (A-B) The distribution of the headings of dendritic (A) and axonal (B) processes in the samples, shown in azimuthal projection. All projections are centered on the average heading direction of the dendritic processes in each sample. Measured directional organization of hippocampal CA1 neuropil reveals approximately perpendicular organization of dendrites and axons. The dendrites are found to run mostly collinearly in the direction perpendicular to the plain of the CA1 cell-body layer. The axons are found to run predominantly within that plane. Additionally, the axons are found to exhibit two distinct directional components: a collinear component and a diffuse component, each containing approximately 50% of all axons. (C) The azimuthal projection plots here show the counts of dendrites and axons moving in different directions in each sample volume. Each heading direction of one dendritic or axonal process can be thought of as a point on a unit sphere. Such direction gets projected onto the unit sphere’s equatorial plane, thus constituting the azimuthal projection plots shown here. Different colors represent the count of the directions projected to the same point, per unit of the unit-sphere’s area.

### 3. Micrometer-scale organization of glia and dendritic spines in hippocampal CA1 neuropil

The random mixing model prescribes that the radial distributions of neuropil components around a reference neural process should not depend on the type of that process, either a dendrite or an axon. This condition, in particular, is met rather well by the measured structure functions for axonal and dendritic neuropil components. For the distribution of dendritic spines (spine-heads) and glia, however, we observe deviations from that rule. In particular, the distribution of dendritic spines is revealed by the respective structure functions to be significantly higher at the proximity of axonal rather than dendritic processes, and for glia we observe the reverse affinity towards dendritic as opposed to axonal surfaces, Figure 5A-B. In the case of glia, more specifically, the average relative density near dendritic processes is measured as 0.17±0.01, while that near axonal surfaces is 0.13±0.005 (Figure 5B). Therefore, we detect the difference in the relative densities of glia near dendritic and axonal processes’ surfaces at p-value <0.001. One can note that spines in neuropil commonly protrude away from dendrites and towards axons, in order to touch the latter to establish synaptic contacts. Thus, the difference observed in the distribution of dendritic spines in Figure 5A may be deemed natural. Similarly, the differences observed in the distribution of glia raise the possibility that certain factors released by dendrites influence or attract glia processes in neuropil towards dendrites.

**Figure 5:**
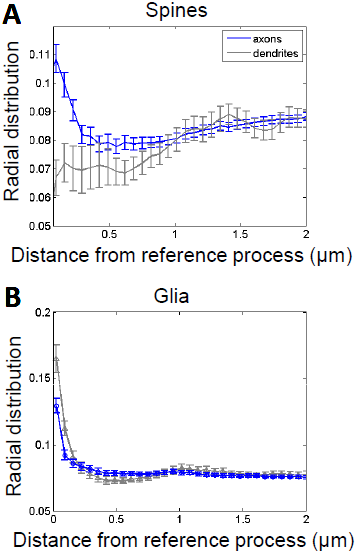
The distributions of glia and dendritic spines in hippocampal CA1 neuropil. (A) The radial distribution functions of dendritic spines relative to axonal and dendritic reference processes. (B) The radial distribution functions of glia relative to axonal and dendritic reference processes. Radial distribution functions of dendritic spines and glia reveal deviations from random mixing model of neuropil. Specifically, the radial distribution functions of dendritic spines reveal significantly higher affinity towards axonal as opposed to dendritic surfaces, and that of glia reveals a reverse affinity towards dendritic as opposed to axonal surfaces. While for dendritic spines the observed affinity can be natural, given the necessity for dendritic spines to protrude away from dendrites and towards axons in order to form synapses, for glia the observed effect may imply presence of novel mechanisms attracting glia towards dendrites. The legend is for all plots. Error bars are the standard error of the mean.

### 4. Distributions of local sizes of neural processes in hippocampal CA1 neuropil

The distribution of sizes of neural processes in neuropil has been a subject of significant attention in neuroanatomy (Braitenberg and Schuz, 1998; Mishchenko et al., 2010; Shepherd and Harris, 1998; Sorra and Harris, 2000). A comprehensive measurement of average neural processes’ sizes had been performed in (Mishchenko et al., 2010). Here, we extend these measurements by characterizing the distribution of neural processes’ local cross-section sizes, that is, such measured along a neural process’ lengths rather than on average. The sizes of axonal and dendritic processes are known to vary significantly even within a single process over the space of just a few micrometers (Braitenberg and Schuz, 1998). Thus, such local size distributions may be important for understanding the local organization of neuropil. We present our measured size distributions in Figure 6. We observe that these distributions are essentially exponential both for axonal and dendritic neuropil components. For spine-heads, the exponential shape of size distribution had been known in the past, and an explanation had been offered based on information-theoretic arguments (Varshney et al., 2006). Here, we observe that exponential shape is characteristic of the local size distributions in all neuropil components, so that a more generic reason may be appropriate. A particularly simple explanation can be offered based on the properties of exponential distributions as the maximum entropy distributions (Jaynes, 2003), implying that the observed size distributions may be the simple outcome of stochastic competition of neural processes for space in dense and constrained neuropil packing (Chklovskii et al., 2002). Although the distribution of dendritic cross-section sizes in Figure 6 appears to show much greater variability than that for axonal cross-sections, it is unclear if that is just the consequence of significantly smaller sample size available in the data for dendrites vs. axons (30-50 dendrites per sample vs. 500-600 axons), or a sign of some real characteristic intrinsic to the dendritic processes.

**Figure 6:**
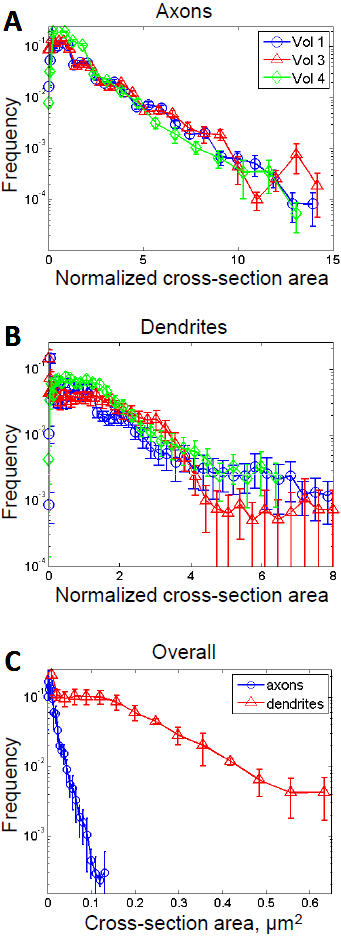
The distribution of axonal and dendritic local cross-section sizes. (A) The distribution of local sizes of axonal cross-sections in sample volumes 1, 3 and 4, normalized to the average axonal cross-section in each volume. The legend shown is for A and B. The error bars are the standard error of the mean. (B) The distribution of local sizes of dendritic cross-sections in sample volumes 1, 3 and 4, normalized to the average dendritic cross-section in each volume. All distributions are well described by the same exponential maximal entropy distribution, suggesting a common mechanism for their origin. (C) The overall distributions of axonal and dendritic cross-section sizes shown on absolute scale; the slope of the distributions is clearly different due to the different size of average axonal (approximately 0.2 p,m) and dendritic (approximately 0.66 p,m) crosssection diameters.

### 5. The relationship between local synaptic connectivity and micrometer-scale organization of neuropil

It had been hypothesized that the small-scale organization of neuropil has direct effect on formation of synaptic connectivity, via influencing the availability of local axonal and dendritic synaptic partners for neural processes in neuropil (Braitenberg and Schuz, 1998; Peters, 1979; Stepanyants and Chklovskii, 2005; Stepanyants et al., 2008). Changes in such organization, therefore, may contribute or be responsible for the disruptions of synaptic connectivity in neural tissues such as observed in many neurodegenerative disorders (Bonda et al., 2010; Hamos et al., 1989; Raff et al., 2002; Scheff et al., 2006; Terry, 2000).

In order to examine this relationship here, we study the variations of the spine density of different dendritic segments in our sample in relation to the structure of the surrounding neuropil, as quantified by measured radial distribution functions. We group all dendritic segments into three categories of high, medium, and low spine-density dendrites, and compare the neuropil structure functions in each such group. Our results are presented in Figure7. We observe that spine-density graded structure functions exhibit no significant variations in the majority of cases, indicating lack of correlations between the spine density of different dendritic segments and local neuropil surrounding, Figure 7B-D. Some features do appear to emerge in Figure 7B-D; however, the deviations that we currently observe are generally one standard deviation or less and therefore cannot be considered significant at this point.

**Figure 7:**
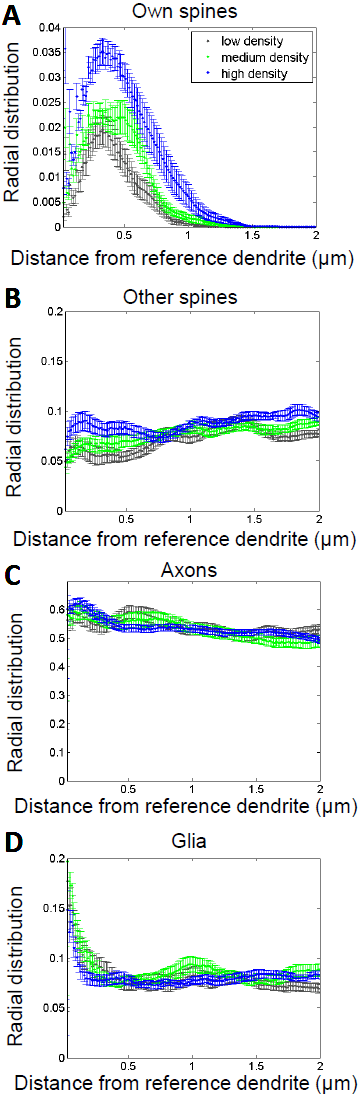
The organization of neuropil around dendritic segments with different spine densities. (A) The radial distributions of reference dendrites’ own spines reveal higher utilization of surrounding neuropil by higher spine-density dendrites. Observed radial distributions indicate that the dendrites of all spine densities place their spines stereotypically in the same zone about 0 to 1 micrometer away, with the maximum spine placement at 400 nm away from the surface of dendritic shafts. (B-D) The other radial distributions exhibit no significant differences in relation to the reference dendrites’ spine-density. Shown here are (B) the radial distributions of the spines of other dendrites, (C) the radial distributions of axons, and (D) the radial distributions of glia, graded by the spine density of the reference dendrite. The low spine-density group was defined as the lowest 50% of spine density dendrites, next 25% comprised the medium spine-density group, and the highest 25% comprised the high-density group. The legend shown is for all graphs. The error bars are the standard error of the mean.

A significant difference is observed in the distribution of the reference dendrites’ own spines, Figure 7A. In this case, the dendrites with higher spine density are clearly observed to exhibit higher densities of own spines in the adjacent neuropil. This may appear simply as a tautological consequence of trifurcating dendrites into low, medium, and high densities; however, this is not so. Since the dendrites with a higher spine-density should place a larger number of spines into their neuropil surrounding, the above result indeed is not unexpected. However, although higher spine density does mean more spines, such spines could be of smaller size leading to similar utilization of neuropil in all spine-density groups, or such spines could have been placed further out from the dendritic shaft, resulting in uniform but more “extended” distributions. Instead, we observe that in all spine-density groups the structure functions have the same shape but different scaling. This can be interpreted in the following way that each spine is created by a dendrite essentially in a “fixed” manner - with similar volume and at similar range of distances from dendritic shaft-just higher density dendrites have more of such spines.

Finally, in Figure 7B we observe that the structure functions of the spines of other dendrites around reference dendrites of all spine-densities are essentially flat, the coefficient of linear regression R=0.0090 μm^−1^ with MSE=0.0031 μm^−1^. We can interpret this indicates the indifference of the spines of other dendrites to the presence of the reference dendrite. That is, the spine zones of nearby dendrites in neuropil appear to interpenetrate or overlap freely and without interactions.

## Discussion

While the ubiquity of anatomical structure in the brain at macro and mesoscales is well known, the structural organization of the brain at the smallest, micrometer scales remained largely unknown. In this work, we aim to fill this gap by examining the structural organization of blocks of neural tissue in s. radiatum of hippocampal area CA1, recently reconstructed using serial section electron microscopy and computerized processing.

Dense reconstructions of neuropil in (Mishchenko, 2009; Mishchenko and Paninski, 2012; Rivera-Alba et al., 2011) offer unique insights into neuropil’s small-scale organization in the sense that they not only provide the reconstructions of a large number of dendritic and axonal processes at nanometer resolution, but also place these in the context of their immediate surrounding in neuropil. To gain insights into the micrometer-scale organization of neuropil here, we calculate the radial distribution functions for neuropil in s. radiatum hippocampal area CA1, earlier reconstructed in the above manner. We analyze obtained measurements statistically and using a modeling approach. Our results indicate that the micrometer-scale organization of neuropil is essentially consistent with disordered packing of axonal and dendritic processes in a tight, space-filling arrangement. At very small distances below 500 nm correlations among neural processes are observed, which are attributed to the physical contact of neural processes in the settings of dense neuropil packing. We also discuss the measurement of the orientational anisotropy of neural tissue and the size distributions of neural processes’ local crosssections in hippocampal CA1 neuropil.

We observe deviations in the micrometer-scale distributions of dendritic spines and glia, whereas dendritic spines are observed to be preferentially located near axonal processes and glia is observed with higher densities near dendritic processes. While for dendritic spines such affinity can be expected, given that spines protrude away from dendrites in order to contact axons to create synapses, there are no known well-established mechanisms that would explain the observed excess of glia near dendrites. The observations here, therefore, may indicate the presence of yet unknown neurobiological factors affecting the development of glia and dendrites in neuropil.

We finally examine the question of the interactions between the local organization of neuropil and synaptic connectivity. While it may be expected that the local organization of neuropil can affect the formation of small-scale synaptic connectivity, we do not observed that to be the case. Indeed, we observe that the synaptic densities of dendritic segments appear to vary completely independently of their surrounding neuropil.

Our analysis relies on characterization of neuropil’s organization using radial distribution functions, here referred to as neuropil structure functions. While these provide a systematic way for assessing the small-scale organization of neuropil, it should be taken into account that structure functions are but one of the possible measures of structure. Specifically, this measure is sensitive to positional coordination, that is, the situations where the presence of one neural process affects the likelihood of encountering other neural processes of specific type in its vicinity. Respectively, our results indicate lack of any micrometer-scale spatial coordination in hippocampal neuropil, but do not imply absence of other types of structure such as depending on neurons’ excitatory/inhibitory type, etc.

It should be also noted that our findings had been produced using a limited sample of hippocampal CA1 neuropil and, therefore, may not apply in other circumstances. Nonetheless, as the past neuroanatomical literature indicates a substantial level of similarity in the organization of neural tissue in different cortical regions (Braitenberg and Schuz, 1998), we may believe that the observations here may be indicative of the organization of the neuropil in mammalian cerebral cortex also more generally.

Our study here focused on a specific question of the structural organization of neuropil viewed as a system of primarily axons and dendrites. Without doubt this focus leaved a large number of similar interesting questions beyond the scope of this paper. A particularly interesting such other question is the structural organization of neuropil around synapses and spines, especially in relation to the types of synapses and other features, or such in respect to the differences between healthy and abnormal tissues. Such topics without doubt will present an interesting venue for future extensions of this work.

## Acknowledgements

The author acknowledges the financial support from Bilim Akademisi - The Science Academy, Turkey, under BAGEP program and the support from the BAP fund - the Fund for Scientific Research Projects, of Toros University, Mersin, Turkey.

